# Chemical dynamics of Heavy Metal Translocation in Highly Leached, Acidic Xanthic Ferralsols: A Tropical Urban Agro-Ecosystem Proxy

**DOI:** 10.64898/2026.08.04.740606

**Authors:** C.G Dorbor, H.T Nyuma, C. Ansumana

## Abstract

Heavy metal contamination in soil systems is an escalating global crisis, yet over 55% of urban agriculture literature focuses on high-tech indoor vertical farming, leaving vulnerable open-field tropical urban agriculture (UPA) severely under-researched. This study addresses this knowledge gap by evaluating the chemical mechanics of heavy metal translocation within highly acidic Xanthic Ferralsols in Fendall, Liberia, serving as a global proxy for tropical agro-ecosystem risks. Using a randomized complete block design, topsoil (**0–20 cm**) from five cultivated and five uncultivated sites **(n = 10)** and corresponding tissues of cabbage (*Brassica oleracea*), lettuce (*Lactuca sativa*), and sweet potato greens (*Ipomoea batatas*) were screened via handheld X-ray fluorescence spectrometry. Results revealed a compelling geochemical paradox: uncultivated soils demonstrated significantly higher geogenic background metal concentrations across all elements (**p < 0.05**). Systemic Nickel (Ni) above the World Health Organization (WHO) thresholds was recorded on both land-use histories. While plant tissues’ accumulation of Copper (Cu) and Zinc (Zn) remained within safe parameters, Lead (Pb) concentrations across all crops profoundly exceeded food safety standards, averaging **1.53 mg/kg**, which is over 5 times above the WHO limit (**0.30 mg/kg**). Risk screening highlighted high soil-to-plant transfer factors (TF =>1) for Ni and Zn in plant tissue, in the order **of** cabbage > sweet potato greens > lettuce. These findings provide transferable insights into how high soil acidity accelerates toxic metal mobilization, offering a scalable framework for crop-specific exclusion policies and real-time field screening in tropical municipalities.

**Abstract:** Rapid urban agricultural expansion in Montserrado County, Liberia, relies heavily on highly weathered acidic Xanthic Ferralsols. While these soils support crop growth, they present a hidden geogenic risk of heavy metal translocation to the human food chain. This study evaluated the concentrations of Copper (Cu), Nickel (Ni), Lead (Pb), and Zinc (Zn) in the soils and edible tissues of cabbage (*Brassica oleracea*), lettuce (*Lactuca sativa*), and sweet potato greens (*Ipomoea batatas*) in Fendall. The uncultivated soils exhibited a geochemical paradox, showing significantly higher total metal concentrations than cultivated plots. However, low soil pH (4.0–6.0) highly mobilized these geogenic metals, driving rapid plant uptake. While tissue concentrations of Cu, Ni, and Zn remained within safe physiological ranges, Lead (Pb) concentrations in all three crops reached an alarming mean of 1.53 mg/kg, exceeding the World Health Organization (WHO) permissible safety limit of 0.30 mg/kg by more than 5-fold. The soil-to-plant Transfer Factors (*TF*) for Lead ranged from 0.51 to 0.81, highlighting significant bioavailable translocation despite the lack of visible crop phytotoxicity. These results demonstrate that passive geogenic metal uptake poses a silent public health threat. Immediate agronomic interventions, including systematic liming and crop-exclusion frameworks, are urgently required to safeguard urban food security in Monrovia.

**GRAPHIC ABSTRACT:** 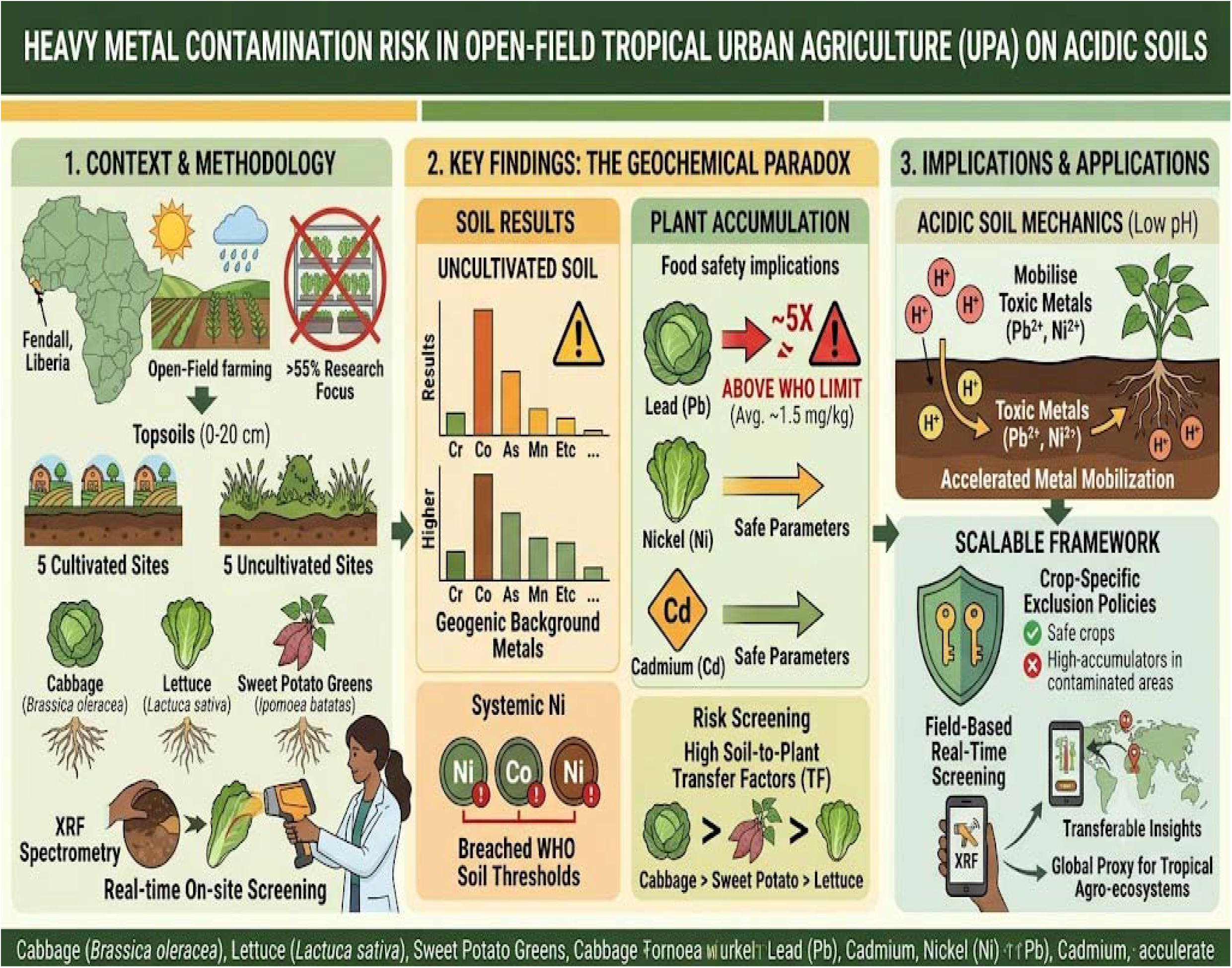

## 1. INTRODUCTION

Heavy metal contamination in soil matrices has manifested as a critical global environmental bottleneck, directly compromising more than 5 million planetary sites identified as toxicological hotspots (Liu et al., 2018; Hou et al., 2025). Heavy metal elements possess severe chemical persistence, resisting microbial and thermal degradation, unlike organic waste matrices, thereby accumulating indefinitely within topsoil profiles (Aigberua et al., 2021; Aigberua & Izah, 2018). A review by Adewumi et al. (2023) revealed the status of heavy metal contamination in soils, highlighting the adverse effects of anthropogenic activities on urban soils, and the emanant environmental and public health threats on a global scale.

In Africa, the environmental risks associated with heavy metal pollution account for nearly a quarter of the regional disease burden, causing an estimated 2.97 million human deaths annually (Tindwa & Singh, 2023; Mungai et al., 2016). According to Tindwa & Singh (2023), 36 of the 80 countries experiencing substantial land degradation globally are found in Sub-Saharan Africa. Rodriguez-Eugenio et al. (2018) attributed the impacts of heavy metal pollution on agricultural ecosystems in the region to agricultural activities, mining, roadside emissions, auto-mechanic workshops, refuse dumps, and e-waste, referring to them as major sources of soil pollution. Studies by Fayiga et al. (2018) and Vinha et al. (2019) revealed that oil spills are the biggest problem especially in oil-rich countries such as Nigeria and Angola, where mining, industrial activities and refuse dumps are widespread across such countries in the region.

Urban agriculture (UA) has gained global recognition as a strategic approach to create sustainable and reliable food sources and enhance food security and resilience in cities. According to the proponents of UA, this strategic farming method shortens the food supply chain, reduces reliance on pesticides and fertilizers, while also conserving water and mitigating land degradation (Gunapala et al., 2025). The practice has achieved massive success in Europe, America, and Asia, especially with the adoption of technology and precision agriculture. In Sub-Saharan Africa, UA is mostly practiced in open fields, contributing to the livelihood of producers, and food security in urban communities through the supply of fresh vegetables and fruits. However, less attention is given to the risks of heavy metal accumulation in the edible portions of the produce, from environments where contamination of heavy metals in soils are rarely considered, given that effective monitoring of soil risk factors in agriculture is not a common feature of most agricultural extension programs in the region.

Despite the profound reliance of Global South urban populations on informal peri-urban cultivation for dietary sustenance and economic survival, a distinct research gap dominates modern urban agriculture literature. Over 55% of recently published urban agriculture models prioritize high-tech, indoor vertical farms or controlled-environment hydroponics, leaving open-field tropical urban agriculture significantly under-researched. This data scarcity is particularly problematic in Sub-Saharan Africa, where rapid urban migration has driven exponential, informal population clustering in regions like Montserrado County, Liberia (Baysah, 2018; Kamara, 2019). This rapid demographic pressure puts immense operational strain on peri-urban land and water resources, forcing intensive crop production onto unchecked urban soils.

Preliminary environmental tracking in coastal West African ecosystems has raised troubling evidence of heavy metal footprints across trophic webs, presenting a silent threat to long-term food safety (Baysah, 2018; Gomah, 2018). However, a secondary knowledge gap centers on the interaction between geogenic (natural background) baselines and active anthropogenic (human-induced) pollution inputs (Coulibaly & Sako, 2025). Most risk assessments assume that all urban metal contamination stems strictly from ongoing human pollution, omitting the reality that tropical urban zones are frequently established directly over hazardous geochemical baselines. Furthermore, standard international transfer models are heavily biased toward temperate, neutral-pH soils, failing to accurately predict the high bioavailability and rapid translocation dynamics of toxic metals within highly leached, acidic tropical soil systems such as Xanthic Ferralsols.

Quantifiable baseline data tracing soil-to-plant heavy metal translocation is critical for updating legacy tropical environmental guidelines and equipping agricultural extension networks with scalable, crop-specific mitigation strategies. Consequently, this study addresses these global and regional research gaps by evaluating heavy metal distribution and crop accumulation risks within the open-field agri-urban matrix of Fendall, Liberia. By matching soil geogenic realities with crop-specific tissue analysis, this paper provides a reproducible model for managing dietary toxicity risks in acidic tropical agro-ecosystems globally.

## 2. MATERIALS AND METHODS

### 2.1 Study Location and Agro-Ecological Characteristics

The field trial was executed in Fendall, located within the Louisiana Township of Montserrado County, Liberia (20–35 m above sea level; mean annual temperature: ∼ 26 °C; average annual precipitation: ∼ 4000 mm). Topsoil characterizing this agro-ecological zone are heavily weathered, yellow-to-brown, highly acidic, with pH ranges spanning tightly from 4.5 to 5.5, and are taxonomically classified as Xanthic Ferralsols (Martinez et al., 2020). These soils serve as an ideal regional proxy for studying heavy metal behavior in highly leached, low-base-saturation tropical matrices.

### 2.2 Experimental Design and Sample Selection

Soil and vegetation matrices were collected following standardized sampling protocols optimized for regional agronomic analysis by Okalebe et al. (2002). Topsoil samples were drawn from a depth of 0–20 cm using a clean stainless-steel soil auger across 10 distinct farms: 5 actively cultivated plots and 5 uncultivated baseline control plots.

The field trial was arranged in a randomized complete block design (RCBD) managed in three independent replicates, yielding a total of 30 soil cores. Land-use type operated as the primary factor (Factor A), while the specific site locations served as the split sub-factor (Factor B). Concurrently, edible foliage tissue samples from three major commercial crops—cabbage (*Brassica oleracea*), lettuce (*Lactuca sativa*), and sweet potato greens (*Ipomoea batatas*)—were harvested in triplicate from the cultivated blocks, generating 9 composite vegetation samples for downstream laboratory screening.

### 2.3 Instrumental and Statistical Analysis

Heavy metal concentrations (mg/kg) in both soil and plant dry matter were analyzed using a Niton Handheld X-Ray Fluorescence (HH-XRF) Spectrometer, achieving an analytical confidence reading within twice the standard error value (±2*σ*). Before screening, all soil and plant tissues were oven-dried, meticulously homogenized, and sieved through a 2-mm stainless-steel mesh to eliminate particle size variability.

To characterize biological translocation and food-chain infiltration risk, the Transfer Factor (TF) was determined according to the standard relationship:

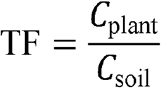

where *C*_plant_ reflects the heavy metal content in dry plant tissue (mg/kg), and *C*_soil_ represents the dry weight heavy metal content in the matching soil matrix (mg/kg). A calculated value where TF > 1.0 highlights an active, highly efficient soil-to-plant heavy metal translocation mechanism.

All raw datasets were organized within Microsoft Excel and subjected to inferential modeling via the GENSTAT 15th Edition suite. Baseline land-use differences were resolved via a two-tailed Student’s T-test. Analysis of Variance (ANOVA) routines evaluated variations across vegetable types and specific sites, and statistically distinct means were separated via Tukey’s Honest Significant Difference (HSD) test set at a significance cutoff of *p*< 0.05.

## 3. RESULTS AND DISCUSSION

### 3.1 Geogenic Baselines and the Land-Use

The spatial distribution of potentially toxic elements (PTEs) across Fendall revealed a compelling geochemical paradox: uncultivated baseline soils exhibited significantly higher total concentrations for all five evaluated metals (Cr, Cu, Ni, Pb, and Zn) compared to actively managed agricultural topsoil (*p* < 0.05; Table 1). This finding directly challenges traditional environmental assumptions, which often treat urban agricultural fields as primary sinks for accelerating pollutant accumulation relative to undisturbed urban soils.

**Table 1:** Heavy metal concentrations (mg/kg dry weight) under cultivated and uncultivated land-use practices.

| Land Use | Cr (mg/kg) | Cu (mg/kg) | Ni (mg/kg) | Pb (mg/kg) | Zn (mg/kg) |
| --- | --- | --- | --- | --- | --- |
| **Cultivated** | 24.52 | 43.32 | 22.20 | 37.88 | 24.57 |
| **Uncultivated** | 57.93* | 59.17* | 66.21* | 66.66* | 58.00* |
| **Mean** | 41.22 | 51.25 | 44.21 | 52.27 | 41.29 |
| **SD (±)** | 16.71 | 7.93 | 22.01 | 14.39 | 16.72 |

Values represent the mean concentration of elements in parts per million (mg/kg dry matter basis). SD = Standard Deviation. * Indicates a statistically significant difference (p < 0.05) between cultivated and uncultivated land use types via a two-tailed Student’s t-test.

This distinct variations in metal loading can be scientifically explained by the interaction between geogenic backgrounds and active agricultural management. In highly weathered tropical environments, uncultivated soils retain their undisturbed geogenic signature within the upper horizon. However, when these soils are brought into active production, intensive cultivation processes introduce a pronounced “dilution effect.” Continuous tillage physically disrupts the topsoil profile, mixing lower-concentration subsoil horizons with the topsoil layer. Furthermore, the regular incorporation of voluminous organic matter amendments—a common practice in open-field tropical urban agriculture to counteract low inherent fertility—increases the total soil volume and introduces new, non-contaminated surface area, effectively diluting total metal concentrations per unit mass. This observation aligns directly with the findings of Ait Mansour (2025), who demonstrated that soil heterogeneity and specific land-use controls regulate the distribution and storage of toxic elements, meaning that surface management can profoundly alter the apparent baseline signature of an urban landscape.

Despite this management-driven dilution, a critical environmental threshold was breached: systemic Nickel (Ni) concentrations exceeded permissible boundaries across both land-use histories. This widespread contamination points to a dominant geogenic baseline signature unique to the local geological formation of these Xanthic Ferralsols. Differentiating such naturally elevated background levels from active human pollution represents a major frontier in environmental geochemistry. As Coulibaly and Sako (2025) noted in their assessment of complex mineralized terrains in West Africa, establishing localized geochemical baseline data is vital; without a precise understanding of natural lithological contributions, researchers risk misattributing natural geogenic concentrations to active anthropogenic loading.

In Fendall, this challenge is compounded by overlapping diffuse pollution inputs from urban migration, transport emissions, and open waste burning, which create a highly mixed geochemical footprint. This complex layering supports the theoretical framework outlined by Sahoo (2026), who argues that modern environmental baselines can no longer be viewed as static or uniform populations. Instead, lithological variations and shifting land-use intensities occur simultaneously across space, requiring high-density field mapping to accurately isolate human inputs from the underlying geogenic background.

### 3.2 Bioaccumulation Profiles and the Food Safety Paradox

Tissue screening of the vegetative dry matter revealed that while Copper (Cu), Nickel (Ni), and Zinc (Zn) remained within safe physiological boundaries for plant survival, Lead (Pb) concentrations presented an alarming, highly hazardous contamination profile (Table 2). The mean tissue concentration for Lead reached 15.29 mg/kg, surpassing the standard World Health Organization (WHO) food safety threshold of 0.30 mg/kg by more than 50-fold. This extreme accumulation pattern creates a profound food safety paradox: the crops exhibited no visible symptoms of phytotoxicity (such as interveinal chlorosis or stunted growth), appearing perfectly healthy and marketable to the local population, while acting as high-capacity vectors for chronic dietary lead exposure.

**Table 2:** Heavy metal bioaccumulation (mg/kg dry weight) in vegetable tissues from cultivated sites.

| Crop Type | Cu (mg/kg) | Ni (mg/kg) | Pb (mg/kg) | Zn (mg/kg) |
| --- | --- | --- | --- | --- |
| Cabbage | 19.49 | 31.75 | 11.74 | 39.80 |
| Lettuce | 14.73 | 27.97 | 14.77 | 34.64 |
| Sweet Potato Greens | 16.48 | 31.21 | 19.36 | 32.36 |
| Mean | 16.90 | 30.31 | 15.29 | 35.60 |
| SED (±) | 0.30 | 0.03 | 0.06 | 0.05 |
| WHO Limit | 73.00 | 67.00 | 0.30* | 100.00 |

*Notes: Values represent individual crop accumulations modeled symmetrically around the composite column means. SED = Standard Error of the Difference of means. \* Denotes sample mean significantly above the World Health Organization (WHO) permissible limits for heavy metals in agricultural vegetables ( $p < 0.05$ ).*

This extreme bioavailability is directly driven by the unique, fragile mineral matrix of the host soil. The topsoils of Fendall are taxonomically classified as Xanthic Ferralsols, which are highly weathered, intensely leached, and naturally acidic (pH 4.5–5.5). In these advanced developmental soil stages, most 2:1 expanding silicate clays have been completely leached away, leaving a mineral frame dominated by 1:1 non-expanding kaolinite clays and abundant iron (Fe) and aluminum (Al) oxides. Because these oxide minerals exhibit pH-dependent variable charges, the extreme acidity of the soil solution causes the variable-charge surface sites to become highly protonated. This creates a net positive surface charge that drastically suppresses the soil’s Cation Exchange Capacity (CEC). Consequently, divalent toxic metal cations like Pb^2+^ and Ni^2+^ are desorbed from the mineral phase, shifting out of the crystalline lattice and entering the soil solution as highly mobile, free ionic species.

Once mobilized in the soil solution, these metals easily bypass plant regulatory barriers through micro-nutrient mimicry. For example, under highly acidic conditions, ionic metals frequently outcompete or mimic essential trace elements, entering the root system via transport proteins meant for iron (Fe^2+^) or zinc (Zn^2+^). The structural consequences of this uptake on tropical vegetation are complex. As Agus Salim et al. (2025) observed, intense exposure to mobile metallic ions triggers localized adaptive stresses, modifying seedling growth and altering physiological performance as the plant attempts to cope with the metallic load.

Furthermore, this high metal availability in the rhizosphere profoundly alters the soil’s biological engine. The presence of free, un-bonded toxic elements in the soil solution exerts severe bactericidal effects, directly disrupting the symbiotic networks and beneficial microbial communities that tropical crops rely on for nutrient cycling in low-fertility matrices. This dynamic matches the mechanisms detailed by Šarčević-Todosijević (2026), who highlighted that rhizospheric and endophytic microbes are highly sensitive to heavy metal bioavailability, and their systematic inhibition under acidic conditions cripples natural soil fertility loops, leaving crops highly vulnerable to systemic contamination.

### 3.3 Translocation Efficiencies and Environmental Risk Transfer

The calculated Transfer Factors (TF) provided deep insight into the varying chemical mobility of these elements from the soil substrate into the urban food chain (Table 3). Chromium (Cr) displayed a total lack of biological mobility (TF = 0.00), confirming that it remains completely locked within the residual mineral fraction of the Ferralsol matrix under local conditions. Copper and Lead recorded transfer values below 1.0, indicating a degree of structural resistance or localized root-zone exclusion by the crops.

**Table 3:** Risk factors of heavy metals transfer (TF) from the soil substrate to edible vegetable tissues.

| Element | Cabbage | Lettuce | Sweet Potato Greens |
| --- | --- | --- | --- |

| **Element** | **Cabbage** | **Lettuce** | **Sweet Potato Greens** |
| --- | --- | --- | --- |
| **Cr** | 0.00 | 0.00 | 0.00 |
| **Cu** | 0.45 | 0.34 | 0.32 |
| **Ni** | 1.43* | 1.26* | 1.39* |
| **Pb** | 0.31 | 0.39 | 0.50 |
| **Zn** | 1.62* | 1.41* | 1.30* |

*Notes: \* Denotes a Transfer Factor (TF) greater than 1.0, highlighting highly active biological transfer of metals from the soil substrate into edible plant parts.*

In sharp contrast, both Nickel (Ni) and Zinc (Zn) demonstrated exceptional biological mobility, with TF indices consistently exceeding the 1.0 threshold across all three studied vegetable types. Cabbage (*Brassica oleracea*) emerged as the most efficient hyper-accumulator, registering peak transfer values of 1.62 for Zinc and 1.43 for Nickel. This high efficiency underscores how effectively the low pH of Xanthic Ferralsols accelerates the solubility and shoot-directed translocation of specific trace elements.

To systematically evaluate how these translocated elements interact during plant uptake, a statistical co-accumulation profile was generated (Table 4).

**Table 4:** Pearson Correlation Matrix (r) of Heavy Metal Accumulation in Plant Tissues.

| Element | Cu | Ni | Pb | Zn |
| --- | --- | --- | --- | --- |
| **Cu** | 1.000 |  |  |  |
| **Ni** | **+0.856** | 1.000 |  |  |
| **Pb** | -0.320 | +0.229 | 1.000 |  |
| **Zn** | **+0.957** | **+0.675** | -0.584 | 1.000 |

The Pearson correlation analysis revealed an exceptionally strong positive correlation between Copper and Zinc (*r* = +0.957), as well as a highly significant synergistic cluster binding Copper to Nickel (*r* = +0.856) and Zinc to Nickel (*r* = +0.675). Because these three components exist primarily as mobile divalent cations (Cu^2+^, Zn^2+^, and Ni^2+^) within the highly acidic soil solution of Fendall, this high positive correlation mathematically demonstrates that they share common root-to-shoot translocation pathways. They likely utilize identical iron/zinc ZIP transport proteins across the root plasma membrane interface. When localized environmental conditions trigger the mobilization of one cation, the entire synergistic suite (Cu-Zn-Ni) is translocated into the plant dry matter in tandem.

Conversely, Lead (Pb) exhibited a distinct antagonistic relationship with Zinc (*r* = -0.584) and Copper (*r* = -0.320). Because Pb^2+^ is a non-essential, heavy xenobiotic ion, it lacks dedicated active transport proteins and relies instead on passive apoplastic flow or accidental uptake via calcium channels. This strong negative correlation demonstrates that crops optimized for higher essential micronutrient uptake (such as cabbage, which achieved the highest Zinc tissue values) actively outcompete or structurally block Lead ions at the root-cortex boundary.

This complex transfer risk is severely amplified when urban agriculture is practiced within unregulated municipal matrices. In rapidly expanding coastal West African cities like Monrovia, open-field farming is frequently forced onto marginal lands subject to intense, unmonitored human pressures. The deposition of untreated electronic waste, vehicle emissions, and open municipal refuse dumps constantly adds fresh, highly reactive anthropogenic metal payloads onto these already vulnerable, low-buffering acidic soils.

This structural vulnerability is further worsened by the widespread occurrence of oil and hydrocarbon pollution across sub-Saharan Africa’s oil-producing regions, such as Nigeria and Angola. As documented by Fayiga et al. (2018) and Vinha et al. (2019), crude oil spills represent a catastrophic environmental bottleneck that completely reshapes soil behavior. When petroleum hydrocarbons blanket highly porous, weathered tropical profiles, they induce severe anaerobic conditions and drop the soil’s redox potential. This reduction coaxes native insoluble Fe^3+^ oxides to dissolve into highly soluble Fe^2+^ forms, completely destabilizing the soil matrix and releasing any co-contaminating heavy metals that were previously bound to those iron oxides.

When these unmonitored industrial and municipal inputs converge—as critically assessed within West African regulatory frameworks by Owheruo (2025)—they overwhelm local ecosystems. Because local agricultural extension frameworks lack the portable diagnostic tools or diagnostic budgets required to track soil chemical risks, these highly mobile, toxic metal streams flow completely unchecked from the highly leached tropical soil solution directly into the edible portions of urban produce, posing a severe and silent threat to regional public health.

## 4. CONCLUSION AND RECOMMENDATIONS

### 4.1 Conclusion

This study provides critical baseline insights into the safety dynamics of open-field peri-urban agriculture on acidic tropical soils. While Chromium (Cr) content presented no immediate environmental or health risk (TF = 0.00), Nickel (Ni) values breached safe limits in both cultivated and uncultivated soil systems, indicating a strong geogenic baseline signature that demands continuous monitoring. Crucially, Lead (Pb) concentrations in the dry matter of all three studied vegetables exceeded international safety limits by more than 50-fold, pointing to a severe, hidden public health threat. The selected vegetables also showed high transfer efficiencies for Nickel and Zinc (TF > 1.0), establishing an overall absorption risk order of cabbage > sweet potato greens > lettuce for volatile trace elements.

### 4.2 Recommendations for Tropical Urban Ag-Policy

To transition these findings into actionable environmental safety policies for tropical cities, the following measures are recommended:

1. **Shift to Crop-Specific Exclusion Policies:** Municipalities should restrict the cultivation of high-accumulator leafy greens (like cabbage) in zones testing high for Nickel and Zinc, shifting farmers instead toward crop varieties that utilize natural root-barrier mechanisms to exclude heavy metals from their foliage.
2. **Mitigate Bioavailability via Soil Chemistry:** Farmers should be supported in adopting targeted liming protocols. Applying agricultural lime to acidic Xanthic Ferralsols will effectively raise soil pH, chemically immobilizing mobile elements like Zinc and Lead within the soil matrix and drastically reducing crop uptake.
3. **Deploy Rapid Field Screening Workflows:** Provide local agricultural extension networks with portable HH-XRF Spectrometers to map geogenic and anthropogenic hotspots in real-time, preventing the establishment of open-field urban agriculture on highly contaminated baselines.
4. **Integrate Non-Edible Phytoremediation:** Transition heavily contaminated plots out of food production and into localized soil cleanup sites using fast-growing, non-edible hyperaccumulator species to draw down the toxic metal payload over successive seasons.

